# Activation of innate immune responses by a CpG oligonucleotide sequence composed entirely of threose nucleic acid

**DOI:** 10.1101/401612

**Authors:** Margaret J. Lange, Donald H. Burke, John C. Chaput

## Abstract

Recent advances in synthetic biology have led to the development of nucleic acid polymers with backbone structures distinct from those found in nature, termed xeno-nucleic acids (XNAs). Several unique properties of XNAs make them attractive as nucleic acid therapeutics, most notably their high resistance to serum nucleases and ability to form Watson-Crick base-pairing with DNA and RNA. The ability of XNAs to induce immune responses has not been investigated. Threose nucleic acid (TNA), a type of XNA, is recalcitrant to nuclease digestion and capable of undergoing Darwinian evolution to produce high affinity aptamers; thus, TNA is an attractive candidate for diverse applications, including nucleic acid therapeutics. Here, we evaluated a TNA oligonucleotide derived from a CpG oligonucleotide sequence known to activate TLR9-dependent immune signaling in B cell lines. We observed a slight induction of relevant mRNA signals, robust B cell line activation, and negligible effects on cellular proliferation.

## Introduction

Chemical modifications have been utilized extensively to improve the binding energy, stability, and tolerability of potential therapeutics, including antisense oligonucleotides, siRNAs, and aptamers, among others. Phosphorothioate modifications in particular have been effective due to their ability to enhance cellular uptake and evade nuclease-mediated degradation in serum. More recently, advances in synthetic biology have led to the development of an expanded set of nucleic acid polymers with backbone structures distinct from those found in nature, termed xeno-nucleic acids (XNAs)(1,2). XNAs are particularly attractive in the field of nucleic acid therapeutics due to several unique properties of several XNAs, mostly notably their high resistance to serum nucleases (2) and their ability to form Watson-Crick base-pairing with DNA and RNA (1). For example, an XNA-based therapeutic for the treatment of neovascular age-related macular degeneration, pegaptanib, was licensed by Eyetech Pharmaceuticals/Pfizer and is currently marketed under the name of Macugen (3,4). Pegaptanib is an XNA-adapted version of an RNA aptamer that binds VEGF165, the misregulation of which can cause abnormal blood vessel growth resulting in bleeding, scarring, and irreversible damage to the photoreceptors(3). 2’-fluoro XNA modifications to pegaptanib were specifically included for evasion of degradation. XNA modifications are under exploration for a variety of other applications as well, including nanostructures for therapeutic delivery of payloads and diagnostics (2,4,5). Thus, XNA-based nucleic acid-based tools hold much promise in the field. However, XNAs have not been extensively evaluated for their ability to induce innate immune responses, and such evaluations will be critical for further therapeutic development of XNAs. Notably, some chemical modifications have been shown to abolish or decrease immune signaling (e.g. locked nucleic acid and 2’-Fluoro substitutions) (6), while others robustly induce immune signaling (e.g. phosphorothioate CpG ODNs). Both phenotypic classes are highly relevant to therapeutics, as some conditions may benefit from stimulation of the immune system (e.g. vaccine adjuvants and anti-cancer applications), while others may benefit from inhibition of immune signaling (e.g.autoimmune disease) or avoidance of immune signaling (e.g. antisense, XNA-mediated payload delivery) (4,5,7–9).

Threose nucleic acids (TNAs) are XNAs in which the natural ribose sugar found in RNA has been replaced by a four-carbon α-L-threofuranose sugar (10). TNA adopts an A-like helical geometry, facilitating stable, anti-parallel Watson-Crick duplexes with complementary strands of DNA and RNA (11,12). Due to a high level of resistance to serum nucleases (13) and recent advances in the use of engineered polymerases that can copy genetic information back and forth between TNA and DNA (1,14), TNA has become an attractive tool for a variety of applications. In fact, *in vitro* selection of TNA aptamers targeting human thrombin (14), HIV reverse transcriptase (15), and ochratoxin A (16) has recently been reported, as well as an antisense TNA targeted to GFP that suppresses GFP expression in live cells (17). Despite recent progress in the utilization of XNAs such as TNA for therapeutic applications, the ability of various XNAs to either avoid or initiate immune responses remains largely unexplored. Here, we set out to evaluate the ability of TNAs to induce innate immune responses in B cell lines by utilizing TNA sequences derived from ligands recognized by human toll-like receptor 9 (TLR9).

TLR9 is a pattern recognition receptor that specifically recognizes DNA. Chemical modifications, specifically phosophorothioate modifications, have long been incorporated into ligands for TLR9, and these ligands have been investigated extensively for a variety of therapeutic applications (e.g. adjuvants, anti-cancer drugs) due to their ability to induce innate immune responses and prime the adaptive immune response. Initiation of innate immune responses through TLR9 by self-DNA (18), unmethylated cytosine-phosphate-guanine (CpG) DNA (19–22), RNA-DNA duplexes (23), and synthetic oligonucleotide sequences containing phosphorothioate-modified CpG oligonucleotide (CpG ODN) motifs (19–22) has been documented. Notably, most of the work to date evaluating TLR9-specific responses has utilized phosphorothioate-modified CpG ODNs (24), demonstrating that TLR9 ligands are amenable to chemical modification while retaining functionality. However, the nature, magnitude, and cell specificity of innate immune induction can differ significantly between the chemically modified phosphorothioate and the natural phosphodiester backbones (25–27), suggesting that there is more to learn about interactions of various natural and unnatural nucleic acids with nucleic acid-sensing receptors.

Currently, there are three classes of CpG ODN. Class A CpG ODNs are characterized by a phosphodiester palindromic motif containing a central CpG motif and a phosphorothioate-modified poly-G string on the 3’ end that has been shown to increase cellular uptake and provide resistance to serum nucleases (20,21,28,29). Class A ODNs induce high levels of interferon (IFN) production in plasmacytoid dendritic cells (pDCs), but only weakly stimulate signaling through TLR9-dependent nuclear factor kappa B (NFκB) activation (21,29). Class B CpG ODNs contain a full phosphorothioate backbone with one or more CpG dinucleotides. Unlike Class A ODNs, Class B ODNs strongly stimulate NFκB signaling, but only weakly stimulate IFN signaling (21,30,31). Class C CpG ODNs are a combination of classes A and B, with a complete phosphorothioate backbone and a CpG-containing palindromic motif, with the capability to induce IFN secretion and activate NFκB signaling (21,22).

For the purposes of this study, we utilized ODN sequence 2395 (5’-tcgtcgtttt<u>cggcgc:gcgccg</u>-3’), one of several commonly studied Class C TLR9 ligands. This ODN has been demonstrated to induce both IFN and proinflammatory responses (21,32,33) and to activate B cells (34). Sequence 2395 was synthesized using a fully phosphorothioate backbone (CpG 2395), a fully phosphodiester backbone (DNA 2395), and a fully TNA backbone (TNA 2395) to enable comparison of the induction of innate immune responses among the three backbones in B cell lines. We then measured induction of various innate mRNAs by quantitative RT-PCR (qRT-PCR), B cell line activation by upregulation of surface-expressed CD86, and proliferative responses. Our data indicate that while TNA 2395 induces only low-level upregulation of IFN and proinflammatory genes at the dosages utilized in this study, TNA 2395 is capable of robust B cell line activation, similar to both CpG 2395 and DNA 2395. While the specific receptor involved in TNA 2395-mediated signaling remains unknown, these data clearly establish that TNA 2395 is capable of inducing innate immune responses in B cell lines.

## Materials and Methods

Unless otherwise noted, all chemicals were purchased from Sigma-Aldrich (St. Louis, MO). Poly I:C was purchased from Invivogen (San Diego, CA). The synthetic TLR9 ligand, CpG ODN 2395, a type C ODN, was purchased from Invivogen (San Diego, CA). DNA 2395 was purchased from Integrated DNA Technologies (Coralville, Iowa). TNA 2395 was synthesized on an Applied Biosystems 3400 DNA synthesizer using standard β-cyanoethyl phosphoramidite chemistry with chemically synthesized TNA phosphoramidites (35). QUANTI-Blue detection reagent for use with the HEK-Blue hTLR9 SEAP reporter cell line was purchased from Invivogen (San Diego, CA).

### Cell Lines

The human B cell lines, Ramos (NIH AIDS Reagent Program, Division of AIDS, NIAID, NIH: Ramos Cells from Drs. Li Wu and Vineet KewalRamani) (36) and Raji (NIH AIDS Reagent Program, Division of AIDS, NIAID, NIH: Raji from Drs. Li Wu and Vineet KewalRamani) (36), were maintained in RPMI medium (Sigma-Aldrich, St. Louis, MO) supplemented with 10% FBS (Sigma-Aldrich, St. Louis, MO) and 2 mM L-glutamine (Gibco, Life Technologies, Grand Island, NY). The human cell line expressing TLR9, HEK-Blue hTLR9 (Invivogen, San Diego, CA), was maintained according to the manufacturer’s instructions. All cell lines were maintained at 37°C in 5% carbon dioxide.

### Cell Stimulatio

For stimulation experiments, Ramos and Raji cells were plated in 1 mL serum-free medium or serum-containing medium at a density of 2 × 10^5^ cells per well in 12 well plates. HEK-Blue hTLR9 cells were plated at 5 × 10^4^ cells per well in 96 well plates. Following plating (Ramos and Raji) or overnight incubation to allow adherence to the plates (HEK-Blue hTLR9), cells were stimulated with the indicated ligands (untreated/vehicle only, poly I:C, CpG ODN 2395, DNA 2395, or TNA 2395) in nuclease-free water at the indicated concentrations.

### RNA Isolation and Quantitatio

For isolation of total cellular RNA, Ramos cells treated with or without ligands for specified TLRs were collected by centrifugation 24 hours post-stimulation and washed twice with 1X PBS, followed by RNA isolation using TRIzol reagent (Invitrogen, Carlsbad, CA) per the manufacturer’s instructions. RNA was precipitated twice and washed with 70% ethanol to remove any contaminants resulting from TRIzol RNA isolation procedures. Isolated RNA was subjected to DNase treatment using Turbo DNase (Ambion, Life Technologies, Grand Island, NY) and quantified on a NanoDrop Spectrophotometer (Thermo Fisher Scientific, Waltham, MA).

### Real Time Quantitative RT-PC

Total cellular was used to synthesize cDNA using random hexamer primers with the ImProm II Reverse Transcription System (Promega, Madison, WI) per the manufacturer’s instructions. The cDNA synthesis was performed using 500 ng RNA. “No RT” and “non-template” (NT) controls were included. Prior to performing Real Time (RT) Quantitative PCR, cellular and viral cDNA, including “no RT” and “NT” controls, were validated by endpoint PCR (40 cycles) using primers specific for 18S rRNA to ensure that “no RT” and “NT” controls were free of contaminating signal. RT-qPCR was performed on each set of cDNA using iTaq Universal SYBR Green Master Mix (Bio-Rad, Hercules, CA) with primers specific to 18S rRNA (forward 5’-CAGCCACCCGAGATTGAGCA-3’ and reverse 5’-TAGTAGCGACGGGCGGTGTG-3’),interleukin 8 (IL-8; forward 5’-CTTCTAGGACAAGAGCCAGGAAGAAACC-3’ and reverse 5’-GTCCAGACAGAGCTGTCTTCCATCAGAA-3’), TNF-alpha (TNFa; forward 5’-GAGTGACAAGCCTGTAGCCCATGTTGTA-3’ and reverse 5’-GCAATGATCCCAAAGTAGACCTGCCCAG-3’), interferon beta (IFNβ; forward 5’-TTGTGCTTCTCCACGACAGC-3’ and reverse 5’-GCACAGTGACTGTCCTCCT-3’), interferon-induced protein with tetratricopeptide repeats 1 (IFIT-1; forward 5’-GTAACTGAAAATCCACAAGACAGAATAG-3’ and reverse 5’-GCAAATGTTCTCCACCTTGTCCA-3’), and 2’-5’-oligoadenylate synthetase 1 (OAS; forward 5’-TGTTTCCGCATGCAAATCAACCATGC-3’ and reverse 5’-GCCAGTCAACTGACCCAGGGCA-3’), according to the manufacturer’s instructions. Each sample was assayed in triplicate to determine technical variability. Relative quantities were determined using the relative quantity (2^-ΔΔCT^) method as previously described (37). Samples were normalized to the specified endogenous control (18S rRNA) to determine expression level relative to total cellular RNA, and these values were plotted relative to the corresponding value for the negative controls (untreated, set to 1). Samples were assayed on the ABI 7500 (Applied Biosystems, Foster City, CA) and analyzed using ABI 7500 Software Version 2.3. Experiments were repeated three times.

### Flow Cytometr

Activation of Ramos and Raji B cell lines was determined by using flow cytometry to detect surface-expressed CD86 after 24 or 72 hours post-stimulation. A phycoerythrin (PE)-labeled antibody for CD86 (B7-2) and a corresponding isotype control, mouse IgG2b kappa, were purchased from eBiosciences (now Thermo Fisher Scientific, Grand Island, NY). Following stimulation of Ramos and Raji cells for 72 hours as described in “cell stimulation,” cells were collected by centrifugation and resuspended in 1X PBS. Cells were incubated with αCD86-PE or isotype control for 30 minutes at 4°C, f°llowed by washing twice in 1X PBS. Cells were then analyzed by flow cytometry using a BD Accuri C6 Flow Cytometer (BD Biosciences, San Jose, CA).

### Proliferation Assay

Proliferation of Ramos cells in response to cell stimulation was assessed by performing cell counts 24 or 72 hours post-stimulation. Cell counts were performed by collecting cells, staining with Trypan blue to exclude non-viable cells, and counting via hemacytometer. Cell counts were confirmed by counting using a BD Accuri C6 Flow Cytometer (BD Biosciences, San Jose, CA).

### Secreted Alkaline Phosphatase (SEAP) Reporter Assay

HEK-Blue hTLR9 cells were seeded at a density of 5 × 104 cells per well in 96 well plates. Cells were stimulated in triplicate with the indicated ligands at the indicated concentrations for either 24 or 72 hours. Following stimulation, QUANTI-Blue reagent was added to a separate 96 well plate according to the manufacturer’s instructions. To assess SEAP activity, 20 µL of supernatant from each well was added to the QUANTI-Blue reagent (180 µL). Reactions were incubated for 2 hours at 37°C and absorbance was detected using a Perkin Elmer Plate Reader at an optical density of 655 nm.

## Results and Discussion

### Differential mRNA upregulation by low and high doses of TLR9 ligands in Ramos Cells

To explore the impact of phosphorothioate, phosphodiester, and TNA backbones on the ability of sequence 2395 to induce innate immune responses, we first sought to determine the responsiveness of the Ramos cells to low and high doses of the previously characterized, commercially available TLR9 ligand, CpG 2395 (phosphorothioate backbone), and the phosphodiester form of CpG 2395, DNA 2395. Cells were also stimulated with the TLR3 ligand, Poly I:C, which is also known to induce innate immune responses in Ramos cells (38), as a positive control. Ramos cells were plated in serum-free medium and stimulated with either 1 µg/mL or 10 µg/mL ligand for 24 hours. Stimulated samples were then subjected to analysis by qRT-PCR to determine RNA levels of a combination of innate immune genes including IFIT-1, IFNβ, IL-8, OAS, and TNFα. These genes were chosen to represent both the interferon response initiated through interferon regulatory factor 7 (IFIT-1, IFNβ, OAS) and the proinflammatory response initiated through NFκB (IL-8 and TNFα) that can be elicited by Class C ODNs, such as CpG 2395. As shown in Figure 1A and B, we observed minimal RNA upregulation with the low dose ligand, while we observed high levels of RNA upregulation with high dose ligands. Notably, the observed response profiles were very similar for CpG 2395 and DNA 2395. These results demonstrate that both the phosphorothioate and phosphodiester forms of sequence 2395 are capable of upregulating interferon and proinflammatory genes upon stimulation in Ramos cells, and that the magnitude of the response is dose-dependent.

**Figure 1.**
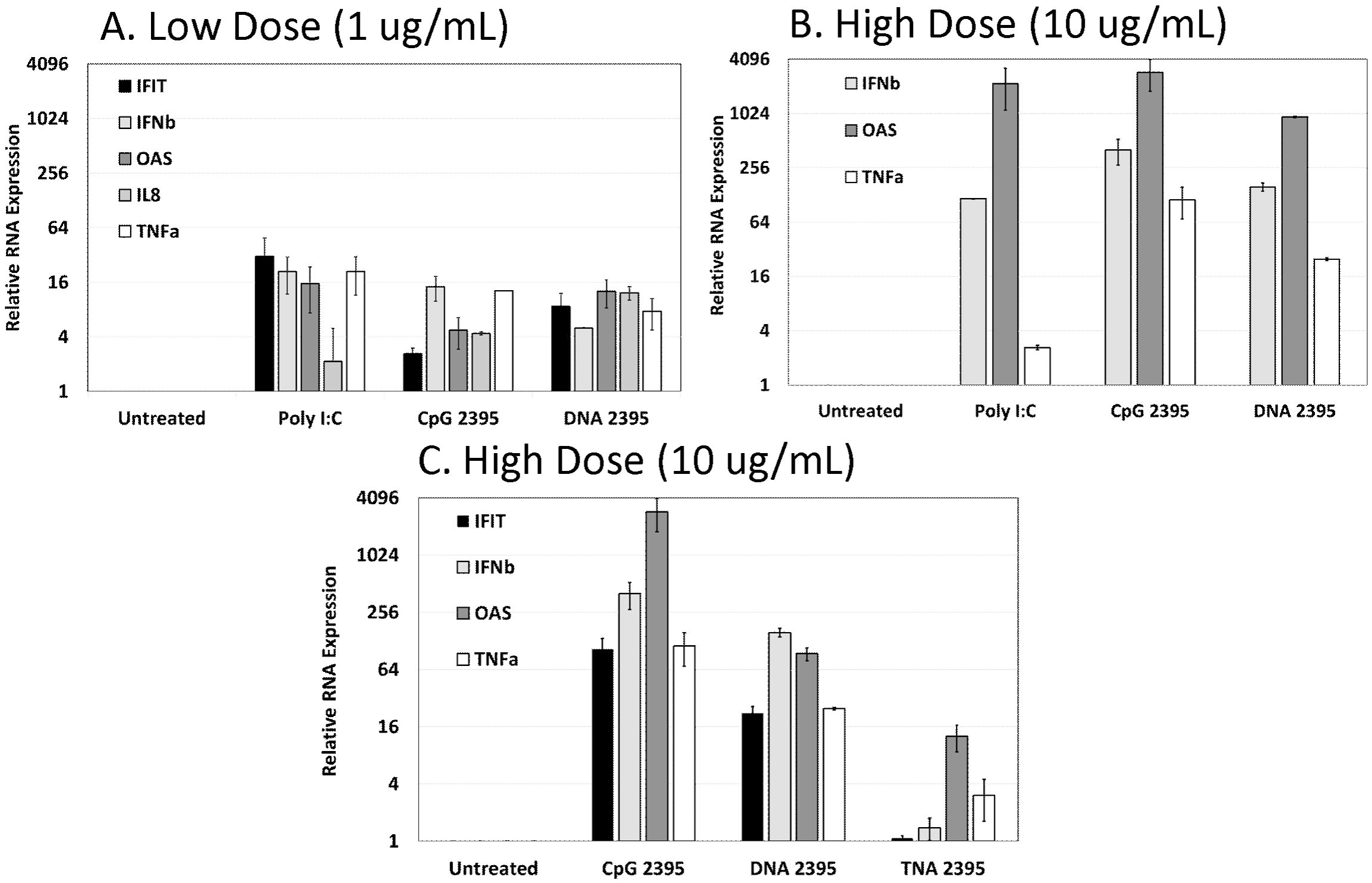
Upregulation of mRNA in response to CpG 2395, DNA 2395, and TNA 2395 in Ramos cells. Ramos cells were plated at a density of 2 × 105 cells in 12-well plates and stimulated with the indicated ligands at the indicated concentrations. After 24 hours, cells were pelleted and washed in 1X PBS, followed by harvest of total cellular RNA. Following DNase treatment of the RNA, cDNA was synthesized and qRT-PCR was performed to examine expression of the indicated genes and 18S rRNA. Relative quantities were determined using the relative quantity (2^-ΔΔCT^) method. Average quantities are shown for technical replicates measured in triplicate. Error bars represent the standard deviation for three technical replicates within each experiment. A representative experiment is shown in each panel. Experiments were repeated three times.

### TNA 2395 displays lower levels of mRNA upregulation in Ramos cells compared to CpG 2395 and DNA 2395

We next wanted to determine whether the 2395 sequence synthesized with a TNA backbone was able to upregulate innate immune genes in the Ramos cells. As shown in Figure 1C, while TNA 2395 was capable of upregulating OAS and TNFα, very little upregulation was observed for IFIT and IFNβ. The low level of IFNβ upregulation was surprising, given the higher level upregulation of OAS, as OAS is an IFN-stimulated gene. It is possible, however, that TNA 2395 induces another type I IFN, such as a subtype of IFNα, which could serve to upregulate OAS. These results demonstrate that while TNA 2395 was capable of eliciting an innate immune response at the level of mRNA via upregulation of interferon (OAS) and proinflammatory (TNFα) genes in the Ramos cells, the magnitude of the response was significantly lower than the levels observed for CpG 2395 and DNA 2395.

### Ramos and Raji cells upregulate CD86 in response to CpG 2395, DNA 2395, and TNA 2395

CpG ODNs are known to trigger both activation and proliferation of primary B cells and various cell lines derived from B cell malignancies (34,39–42). Therefore, we next set out to determine whether stimulation of Ramos and Raji B cell lines by CpG 2395, DNA 2395, and TNA 2395 led to upregulation of the well-known B cell activation marker, CD86 (28,42,43). Ramos cells were stimulated with ligands ranging from 1 µg/mL to 10 µg/mL. After 72 hours, cells were collected and analyzed for CD86 expression by flow cytometry. As shown in Figure 1A, Ramos cells were robustly activated by all three 2395 sequences. The highest ligand dose utilized yielded 4.5-fold induction by CpG 2395, and a 6.5-fold induction for both DNA 2395 and TNA 2395. Interestingly, although TNA 2395 induced only low-level upregulation of innate immune-associated mRNAs explored in the context of this study, activation of Ramos cells via upregulation of CD86 expression by TNA 2395 was similar to that of both CpG 2395 and DNA 2395, indicating that TNA 2395 is able to robustly activate Ramos cells.

In addition to activation of the Ramos cells, the 2395 sequences were analyzed for their ability to activate Raji cells. Notably, Raji cells display a higher level of CD86 expression in their unstimulated state, presumably due to the chronic Epstein Barr Virus infection of the cell line (44). Similarly to the Ramos cells, CpG 2395, DNA 2395, and TNA 2395 were all able to upregulate CD86 expression in the Raji cells above the background level after 72 hours stimulation, albeit to a lesser extent (3-fold) (Figure 2B).

**Figure 2.**
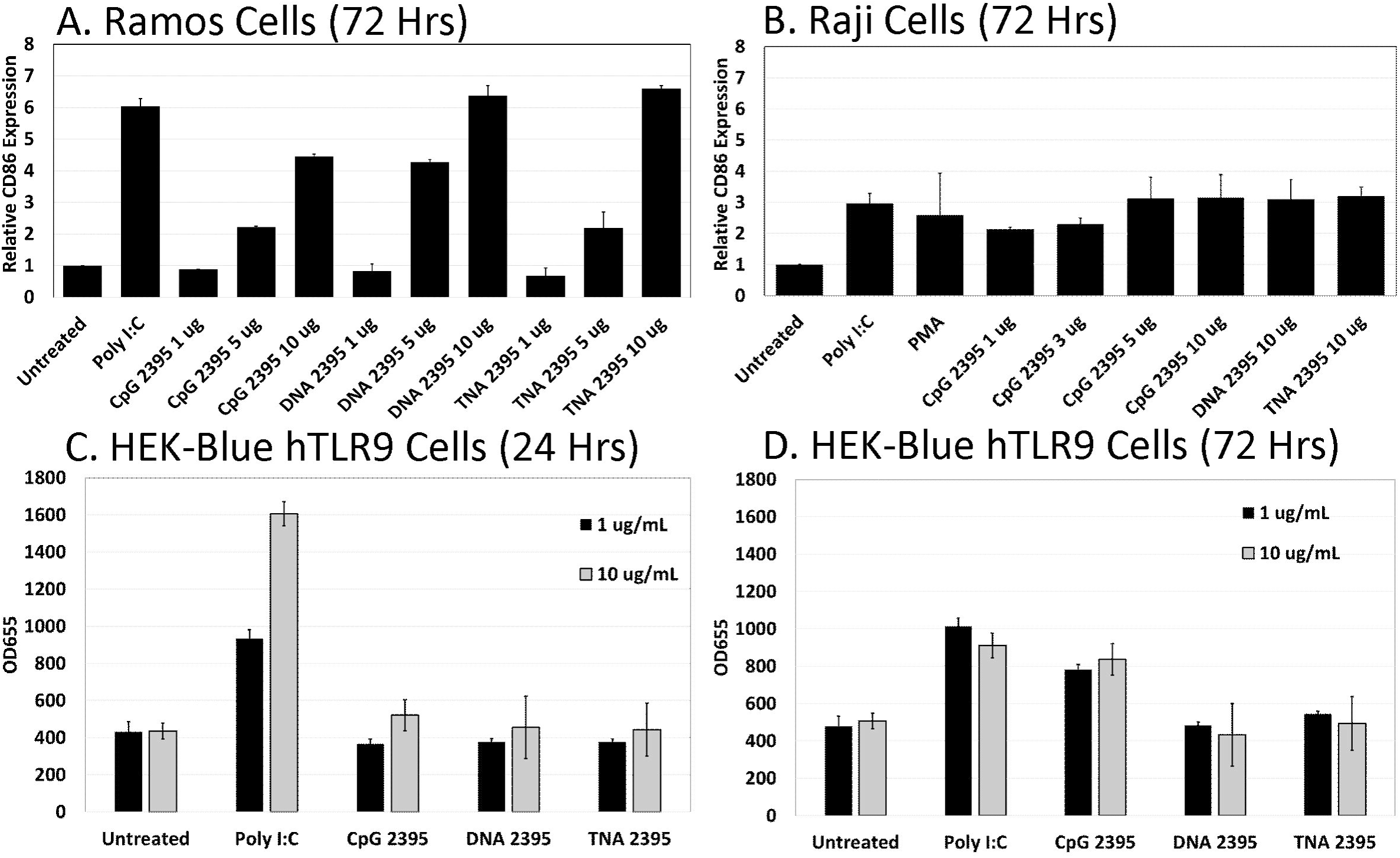
CpG 2395, DNA 2395, and TNA 2395 activate Ramos and Raji cells, while only CpG 2395 induces SEAP reporter activity in HEK-Blue hTLR9 cells. (A and B) Ramos and Raji cells were plated at a density of 2 × 105 cells in 12-well plates and stimulated with the indicated ligands at the indicated concentrations. After 72 hours, cells were harvested for flow cytometric analysis of surface-expressed CD86 expressing using a PE-labeled antibody specific for CD86. A PE-labeled isotype antibody was included as a control. Data are shown relative to the untreated control for Ramos (A) and Raji (B) cells. Error bars represent the standard deviation between two replicates within a single experiment. A representative experiment is shown in each panel. Experiments were performed three times. (C and D) HEK-Blue hTLR9 cells were plated at a density of 5 × 104 in 96 well plates and allowed to adhere overnight. Cells were then stimulated with the indicated ligands at the indicated concentrations. After 24 (A) or 72 (B) hours, supernatants were harvested and tested for SEAP activity using QUANTI-Blue reagent. Samples were assayed in triplicate. Error bars represent standard deviations for three replicates within a single experiment. A representative experiment is shown in each panel. Experiments were performed three times.

**Figure 3.**
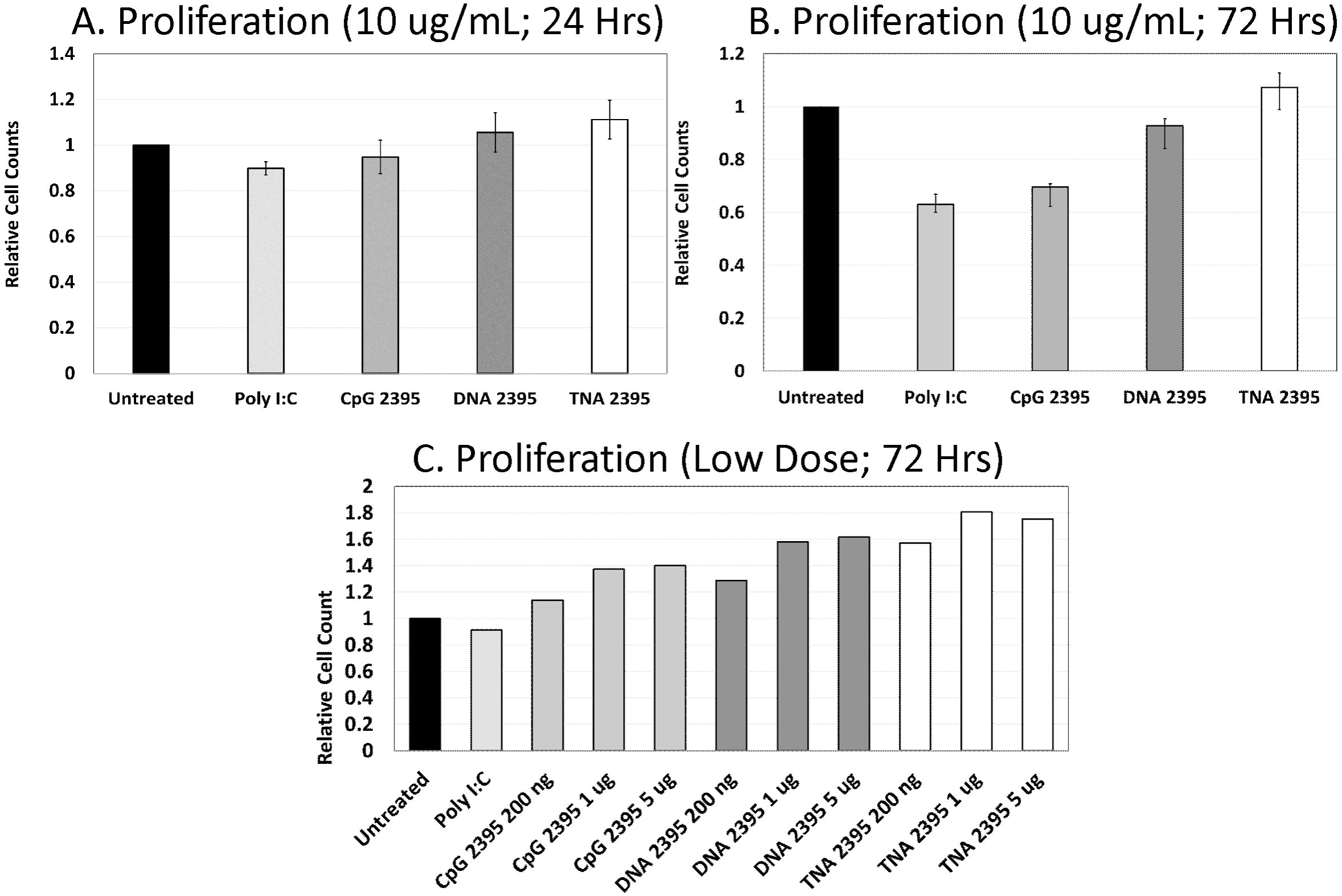
CpG 2395 decreases B cell proliferation, while DNA 2395 and TNA 2395 do not alter B cell proliferation. Ramos cells were plated at a density of 2 × 105 cells in 12 well plates and stimulated with the indicated ligands at the indicated concentrations. After 24 (A) or 72 (B and C) hours, cells were collected and counted using a hemacytometer. Cell counts were confirmed using a flow cytometer to ensure counting accuracy. Cell counts relative to the untreated control are shown. Experiments were repeated three times.

Together, these results confirm prior studies demonstrating that CpG ODNs containing phosphorothioate and phosphodiester backbones are able to activate B cell lines. In addition, these data also show that a TNA backbone is able to robustly activate B cell lines, despite only low-level upregulation of innate immune-associated genes examined in this study as compared to CpG 2395 and DNA 2395.

### HEK-Blue hTLR9 cells respond to CpG 2395, but fail to respond to DNA 2395 and TNA 2395

HEK-Blue hTLR9 cells are a commercially available cell line specifically designed to analyze the immunological properties of TLR9-specific ligands. These cells stably express secreted alkaline phosphatase (SEAP) under control of the NFκB promoter. Originally, we planned to utilize a lentiviral vector expressing a TLR9-specific guide RNA and CRISPR/Cas9 to knockout TLR9 in the HEK-Blue hTLR9 cells to determine whether TNA 2395 was specifically signaling through TLR9, as B cell lines are notoriously difficult to transduce. However, as shown in Figure 2C and 2D, while the HEK-Blue hTLR9 cells respond to Poly I:C after 24 and 72 hours of stimulation and to CpG 2395 after 72 hours of stimulation, they fail to respond to either DNA 2395 or TNA 2395. Thus, we were unable to examine the ability of TNA 2395 to specifically stimulate TLR9 within the scope of this study. Interestingly, a recent report demonstrated that expression of human macrophage scavenger receptor 1 gene (hMSR1), which is not expressed in the HEK-Blue hTLR9 cells, was necessary for induction of SEAP by CpG ODN containing a phosphodiester backbone in the HEK-Blue hTLR9 cells (45). When hMSR1 was transfected into the HEK-Blue hTLR9 cells, uptake and SEAP activity of CpG DNA was shown to be equivalent to that of CpG ODN with a phosphorothioate backbone (45). Thus, it is possible that addition of hMSR1 to our HEK-Blue hTLR9 cells would allow uptake of DNA 2395 and TNA 2395, conveying the ability of the cells to respond to these ligands.

### CpG 2395 and DNA 2395 inhibit Ramos cell proliferation at high doses, but stimulate proliferation at low doses

In addition to activation of B cells, CpG ODNs have been shown to induce proliferation of primary B cells and some cell lines derived from B cell malignancies (28,34,39,41,43). However, stimulation with CpG ODN has also been demonstrated to induce apoptosis in other B cell lines (9,41). Therefore, we next sought to determine whether stimulation of Ramos cells by CpG 2395, DNA 2395, and TNA 2395 leads to a proliferative response. We examined proliferative responses by cell counting at both 24 and 72 hours at a ligand dose of 10 µg/mL. At 24 hours post-stimulation, we did not observe a difference in cell counts. This was expected, as the cells would not have undergone many rounds of cell division. At 72 hours, we observed a reduction in cell number compared to the untreated control for Poly I:C and CpG 2395. Poly I:C is a well-known inducer of apoptosis through TLR3-mediated signaling, so the observed decrease was expected. CpG ODNs have specifically been shown to induce apoptosis in Burkitt’s Lymphoma-derived cell lines, such as the Ramos cells (46). DNA 2395 demonstrated a slight reduction in cell counts at 72 hours, but not to the magnitude of Poly I:C nor CpG 2395. TNA 2395 cell numbers remained equivalent to the untreated control, suggesting that TNA 2395 does not alter proliferative responses despite robust activation of B cell lines, similarly to responses observed for ligation of TLR7 in B cells (40).

In conclusion, the data presented here demonstrate that similar to CpG 2395 and DNA 2395, the 2395 sequence composed entirely of TNA is able to stimulate some innate immune responses. Notably, the ability of TNAs to induce immune responses has not previously been examined. Similar to sequences composed of phosphorothioate backbones, the TNA equivalents are highly resistant to serum nucleases (13), making them an attractive tool for the future development of nucleic acid-based therapeutics. Recent advances in TNA synthesis using engineered polymerases have allowed for the selection of TNAs that target human thrombin (14) and HIV reverse transcriptase, opening the door for selections against a variety of other protein targets. In addition, a recent study demonstrated that antisense TNA oligonucelotides are able to effectively suppress GFP expression in living cells (17). While a potential benefit of chemically modified nucleic acids is the ability to evade the immune response (e.g. antisense applications), chemically modified nucleic acid therapeutics such as TNAs could also be highly useful as stimulators of immune responses (e.g. adjuvants) or inhibitors of immune responses (e.g. anti-inflammatory therapies). Thus, further exploration of the utility of TNAs and other chemical modifications are needed, including their ability to stimulate or inhibit immune responses and the specificity of these responses.

Whether TNA 2395 signals specifically through TLR9, as with CpG 2395 and DNA 2395, remains to be determined. TNA is capable of antiparallel, Watson-Crick base-pairing with complementary DNA, RNA, and TNA oligonucleotides, and the self-paired TNA/TNA duplex occurs through an A-like helical geometry (10,47–49). Thus, it is possible that TNA 2395 could also signal through endosomal TLRs that recognize RNA, such as TLRs 3, 7, or 8, or even a variety of other nucleic acid-sensing receptors should the TNA escape the endosome. Further experiments beyond the scope of this study will be required to determine the specificity of TNA-induced signaling.

